# Co-transcriptional capping using an RNA capping enzyme-T7 RNA polymerase fusion protein

**DOI:** 10.1101/2023.10.28.564488

**Authors:** S. Hong Chan, Louise Tirole, Daniel Kneller, Theresa M. Kelley, Nan Dai, G. Brett Robb

## Abstract

mRNA vaccines and therapeutics are highly effective and can be developed and manufactured with a relatively short lead time. Here we report an all-enzyme platform to generate capped synthetic RNA in a one-step process based on an RNA capping enzyme-T7 RNA polymerase fusion protein. Under standard in vitro transcription reaction conditions, the fusion protein, in conjunction with an RNA cap 2′-O-methyltransferase, can generate synthetic mRNA with up to 95% of Cap-1 incorporation, greatly simplifying mRNA manufacturing workflows.

## Main

mRNA vaccines advanced from experimental to clinical in less than a yearto address the COVID19 pandemic ^1^. The successful rapid manufacturing of the mRNA vaccines against SARS-CoV-2 is the cumulation of decades of research on virus biology, immunology, DNA and RNA sequencing technologies, in vitro transcription, and lipid nanoparticle technologies. Yet, to meet the foreseeable future demand of mRNA vaccines and therapeutics (such as cancer immunotherapies) improved mRNA manufacturing technologies are needed.

The 5′ Cap-1 structure is one of the critical quality attributes (CQAs) of synthetic mRNA ^2,3^ due to its important biological roles in efficient protein translation and evading innate immune responses ^4^.

Currently, two methods of Cap-1 incorporation are being used in mRNA therapeutics manufacturing: post-transcriptional capping using a combination of a trifunctional RNA capping enzyme with vaccinia cap 2′-O-methyltransferase ^5–7^ and co-transcriptional capping using chemically synthesized capped dinucleotides ^8,9^. Each method offers advantages and tradeoffs in terms of complexity of manufacturing and cost ^10^. A highly simple and cost effective one-pot Cap-1 mRNA synthesis can further enable development and implementation of mRNA therapeutics.

To that end, we first investigated whether including RNA capping enzymes in T7 RNAP IVT reactions could achieve satisfactory level of Cap-0 incorporation. RNA polymerase and RNA capping enzyme reactions can be biochemically incompatible with each other. For example, vaccinia RNA capping enzyme (VCE) has been shown to hydrolyze free ATP and GTP ^11–14^, building blocks for in vitro transcription. The high magnesium ion concentration used in high yield IVT conditions can inhibit RNA capping enzymes ^7.^ Inorganic pyrophosphate generated by T7 RNAP at each nucleotide incorporation can promote the reverse guanylylation activity of RNA capping enzymes and reduce the yield of capped RNA products ^12^. Typical IVT reactions include an inorganic pyrophosphatase to hydrolyze inorganic pyrophosphate to inorganic phosphate to reduce pyrophosphorolysis of the RNA polymerase ^15.^. It is unclear if a concentration of inorganic pyrophosphate inhibitory to RNA capping enzymes will accumulate as IVT proceeds. The hydrolysis of inorganic pyrophosphate may further contribute to increased levels of free magnesium ions that can inhibit capping enzyme activity ^7^. Using a hRNase 4-LC-MS-based cap analysis method ^16^, we found that 90% or higher level of Cap-0 incorporation is only achievable using 500 nM of RNA capping enzyme at 45°C (Supporting data 1). Such a relatively high enzyme concentration would poseoperational and cost challenges in implementation.

We therefore generated a fusion protein that links a recently characterized single subunit Faustovirus RNA capping enzyme (FCE) ^7^ to T7 RNA polymerase ^17^. We first verified that the FCE::T7RNAP fusion was highly active in RNA capping under standard capping conditions (Supporting data 2). To help understand the effect of 5′ sequence to enzymatic capping and IVT yield, we generated a panel of IVT templates encoding transcripts with different 5′ UTRs with shared ORF (firefly luciferase), 3′ UTR (based on a publicly available Comirnaty sequence ^18^) and a poly(A) tail (A60). We found that the FCE::T7RNAP fusion protein generated up to 90% Cap-0 RNA for the transcripts containing the human hemoglobin beta subunit (HBB) or pRNA21 from linearized plasmid (Fig. 1b and Supporting data 3) or PCR-amplified DNA templates (Supporting data 4) at 37°C under standard IVT conditions supplemented with SAM (co-substrate required for methyltransferase activity), demonstrating that the FCE::T7RNAP fusion enzyme can generate Cap-0 RNA in a one-pot, one-step all-enzymatic reaction. On the other hand, only ∼30% of the FLuc 5′UTR transcript was capped under these co-transcriptional capping conditions. For a challenging transcript like the FLuc 5′UTR transcript, we found that the level of Cap-0 incorporation could be improved by increasing the reaction temperature (Fig. 1c, Supporting Data 5) and/or increasing the concentration of FCE::T7RNAP fusion used in the reaction (Fig. 1d, Supporting Data 5).

**Figure 1.**
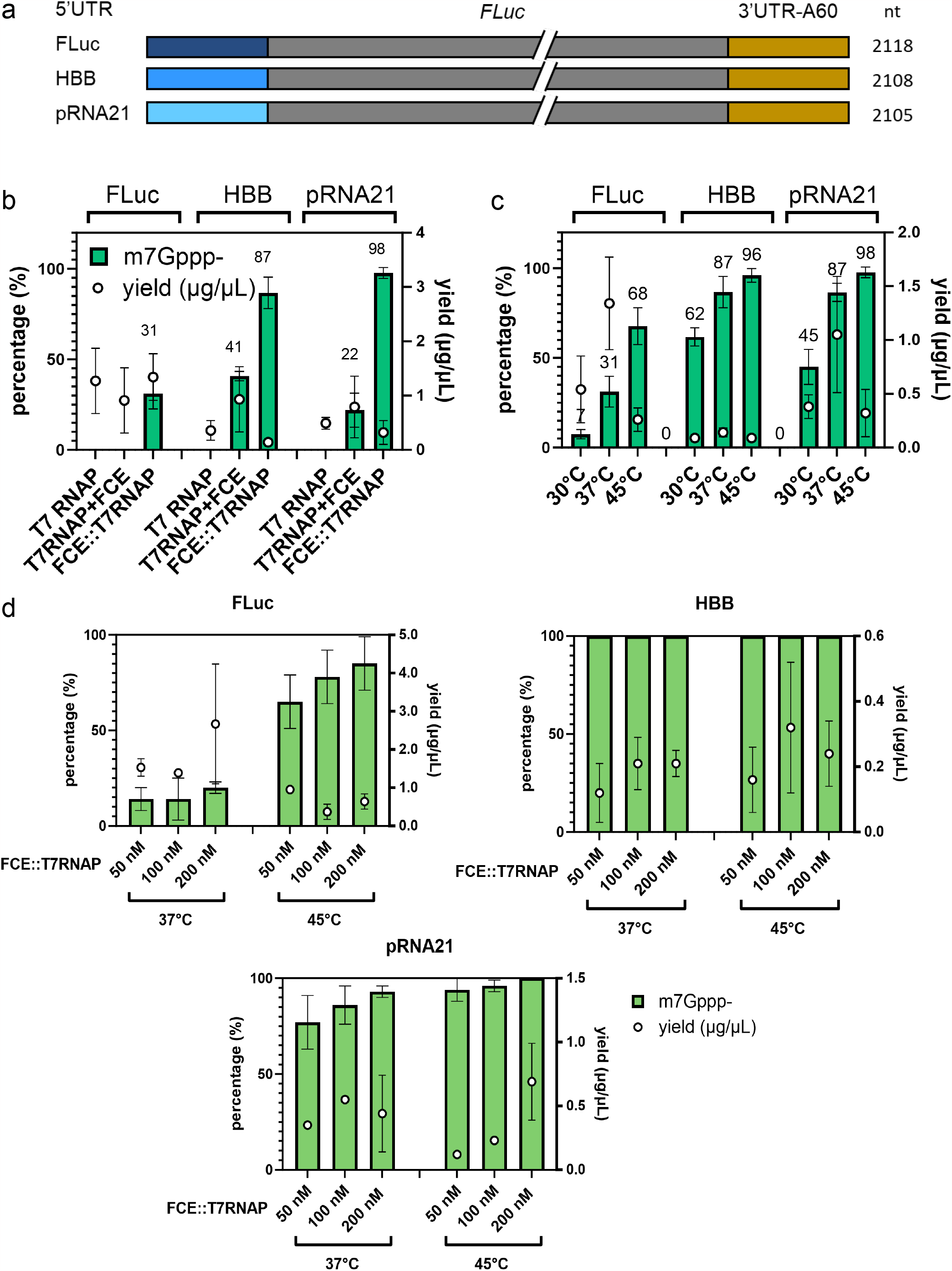
(a) Design of the 5′UTR panel IVT templates. The three transcripts shared the same gene body (firefly luciferase), a 3′UTR derived from the publicly available sequence of Comirnaty, followed by a poly(A) tail of 60 adenosines. (b) Comparison of Cap-0 incorporation and yield of the three 5′UTR transcript using T7 RNAP (80 nM), one-pot T7 RNAP and FCE (100 nM each), and the FCE::T7RNAP fusion (100 nM) at 37°C. No Cap-0 transcript was detected for FLuc using a mixture of T7 RNAP and FCE. Approximately 31% of Cap-0 FLuc 5′UTR transcript was generated using the FCE::T7RNAP fusion. The HBB 5′UTR transcript was capped up to 41% or 87% using the T7 RNAP/FCE mix, or the FCE::T7RNAP, respectively. The pRNA21 5′UTR transcript was capped up to 98% using the FCE::T7RNAP fusion, while only 22% of the transcripts were capped using the T7 RNAP/FCE mix. The yield of about 1.5 μg/μL was achieved for the FLuc 5′UTR transcript using T7 RNAP, the T7RNA/FCE mix of the FCE::T7RNAP fusion. The yield of the HBB and pRNA21 5′UTR transcripts were lower at ∼0.5 μg/μL using T7 RNAP, ∼1 μg/uL using the T7RNAP/FCE mix and ∼0.2 μg/uL using the FCE::T7RNAP fusion. A detailed breakdown of enzymatic capping intermediates can be found in Supporting data 3. (c) Increasing reaction temperature from 30°C to 45°C improves Cap-0 incorporation. Except the HBB 5′UTR transcript whose yield remained low at 0.1-0.2 μg/uL, we observed an increase in yield going from 30°C to 37°C but a drop going from 37°C to 45°C. 100 nM of FCE::T7RNAP was used. A detailed breakdown of enzymatic capping intermediates can be found in Supporting data 3. (d) Increasing FCE::T7RNAP concentration improved Cap-0 incorporation. The FLuc 5′UTR transcript, being a challenging transcript to cap under co-transcription conditions using the FCE::T7RNAP fusion, can be capped to a higher level by using more FCE::T7RNAP and increasing the reaction temperature to 45°C. The pRNA 5′UTR saw improvement in Cap-0 incorporation that approached 100% as the FCE::T7RNAP concentration was increased to 200 nM. The HBB 5′UTR transcript could be capped up to 100% using 50 nM FCE::T7RNAP fusion. A detailed breakdown of enzymatic capping intermediates can be found in Supporting data 5.

The fact that the FLuc 5′UTR was challenging to cap by the FCE::T7RNAP fusion is interesting because the same transcript was capped to 94-100% using the two-step IVT and enzymatic capping workflow (Supporting data 6). The low co-transcriptional capping activity but high post-transcriptional level suggests that the transcription kinetics for the FLuc 5′UTR sequence may play a role in the level of co-transcriptional capping.

When we focused on transcription, we noticed that the three 5′UTR contributes to different levels of yield by T7RNAP alone (Fig. 1b), pointing to the less-understood role of 5′ sequence in IVT. The yield of the HBB 5′UTR transcripts was particularly low when T7RNAP or FCE::T7RNAP was used (∼0.1 μg/μL), probably due to the use of T7 promoter phi2.5 to accommodate adenosine as initiating nucleotide (Fig. 1b-d). More research will be needed to pinpoint factors such as the properties of T7 phi2.5 promoter, initiating RNA sequences, and the propensity of 5′UTR to abortive initiation^19–21^. As far as mRNA synthesis is concerned, transcription yield can be improved by modififying reaction conditions such as higher template concentrations (Supporting data 7)^22,23^ and through process engineering.

Synthetic mRNA intended for therapeutic applications requires the Cap-1 structure to help evade innate response^4,24–26^. It has been shown that vaccinia cap 2′-O-methyltransferase (2′OMTase) is compatible with FCE and VCE in a one-pot format to install the Cap-1 structure on synthetic RNA ^7^. Here, we show that vaccinia 2′OMTase can be used in FCE::T7RNAP co-transcriptional capping to generate >95% Cap-1 RNA from the HBB and pRNA21 5′UTR DNA templates (Fig. 2a, Supporting Data 8). We further evaluate if the FCE::T7RNAP fusion enzyme can generate longer Cap-1 RNA. We synthesized a DNA template based on the publicly available sequence of the Moderna COVID-19 vaccine mRNA-1273 with modifications (Supporting data). We showed that the FCE::T7RNAP fusion can generate up to 90% Cap-1 mRNA-1273* molecules at 37°C and 45°C reaction temperatures (Fig. 2b, Supporting Data 9). The yield of transcription was ∼0.5 μg/μL, approximately 50% lower compared to T7 RNAP under the same conditions (Fig. 2b).

**Figure 2.**
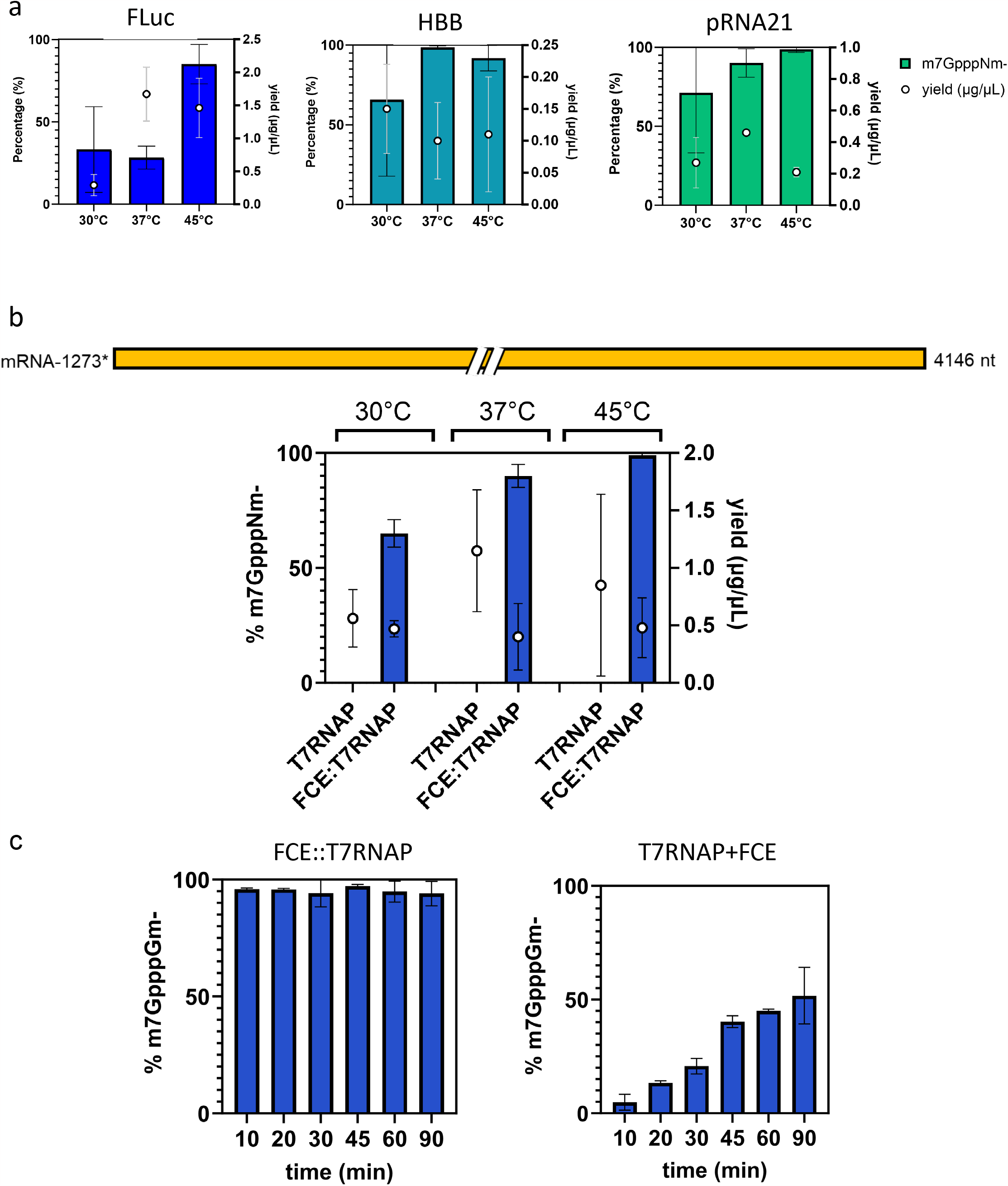
(a) One-pot, one-step Cap-1 RNA synthesis. 200 nM of the FCE::T7RNAP fusion and 5U/μL of vaccinia cap 2′-O-methyltransferase was used to synthesize capped RNA for the 5′UTR panel. The level of Cap-1 incorporation increased as the reaction temperature increased. Increasing the reaction temperature benefited the FLuc 5′UTR most by bringing Cap-1 incorporation from ∼30% at 37°C to ∼90% at 45°C. The HBB and pRNA21 5′UTR achieved >90% Cap-1 incorporation at a yield of 0.1 μg/μL (HBB 5′UTR) and 0.2-0.4 μg/μL (pRNA21 5′UTR) at 37°C or 45°C. A detailed breakdown of enzymatic capping intermediates can be found in Supporting data 8. (b) The FCE::T7RNAP fusion can be used to generate a 4 kb RNA with high levels of Cap-1 incorporation. A synthetic construct based on the Moderna mRNA-1273 sequence (∼4.1 kb) was synthesized using 200 nM of the FCE::T7RNAP fusion or 80 nM of T7 RNAP at 30°C, 37°C or 45°C. Although the transcription yield at 37°C and 45°C was approximately 50% of the level achieved by T7 RNAP (0.5 μg/μL compared to 1 μg/μL), ∼95% or ∼98% of Cap-1 RNA was synthesized at 37°C or 45°C, respectively, demonstrating the applicability of the FCE::T7RNAP fusion in synthesizing highly capped RNA in a co-transcriptional format. Datapoints are averaged values of three independent experiments. Error bars represent one standard deviation. A detailed breakdown of enzymatic capping intermediates can be found in Supporting data 9. (c) 5′ capping and transcription take place in cis. A time course from 10 to 90 minutes showed that the pRNA 5′UTR transcript was capped to ∼95% at 10 minutes using 100 nM of the FCE::T7RNAP fusion and 5 U/μL vaccinia cap 2′-O-methyltransfease. The level of Cap-1 incorporation maintained at the same level up to 90 minutes. Control experiment using 80 nM T7 RNAP, 100 nM FCE and 5 U/μL of vaccinia cap 2′-O-methyltransfease resulted in Cap-1 incorporation from 0 to ∼50% over 90 minutes. Datapoints are averaged values of two independent quadruplicated experiments. Error bars represent one standard deviation.

Finally, to investigate whether 5′ capping and transcription take place in cis, i.e., the 5′ end was capped by the same FCE::T7RNAP fusion molecule that transcribed the RNA, we conducted a time-course experiment using the pRNA21 5′UTR template (Fig. 2c). (Fig. 2c). We reasoned that if 5′ capping and transcription take place in cis, we would achieve a high level of cap incorporation at early time points as in later time points. In the presence of 5 U/μL vaccinia 2′-O-methyltransfease, RNA generated by 100 nM of the FCE::T7RNAP fusion maintained >95% Cap-1 incorporation starting from the earliest measured timepoint (10 minutes) up to 90 minutes. In contrast, an equimolar mixture of T7 RNAP and FCE (100 nM each) saw a gradual increase in Cap-1 incorporation up to only ∼50% at 90 minutes. With an understanding that vaccinia cap 2′-O-methyltransferase is highly efficient at converting Cap-0 to Cap-1 (Supporting Data 8-9), the observation that a high cap incorporation level was maintained from an early time point up to 90 minutes suggests that 5′ capping occurs primarily in cis - through an intermolecular process in the FCE::T7RNAP fusion. Quantitation of the transcript showed that the FCE::T7RNAP fusion was generating transcript steadily over time (Supporting Data 10).

In this study, we showed that an FCE::T7RNAP fusion can effectively generate Cap-1 transcripts from DNA templates in a one-pot, one-step co-transcriptional format under typical IVT conditions. However, we did not optimize conditions to try to achieve higher yield. We observed variability of transcription yield by the fusion enzyme, especially compared to using T7 RNAP alone. It has been known in the mRNA manufacturing sector that although all mRNA synthesis processes use the same set of reaction reagents, reaction conditions such as concentration of NTPs, template, enzyme, incubation time, and reaction temperature need to be fine-tuned for individual RNA sequences and T7 RNAP variants to maximize yield and achieve high quality RNA synthesis (such as minimizing dsRNA formation). Factors such as sequence design of the DNA template can also impact the outcome of IVT. For example, we showed that the capping performance and transcription yield of FCE::T7RNAP was impacted by 5′UTR sequence. Although research has been done to help illuminate the role of 5′UTR in protein translation in vivo ^27,28^, more research is needed to help understand the role of 5′ sequence in IVT and enzymatic capping.

The FCE::T7RNAP fusion enzyme produced >90% of Cap-1 incorporated mRNA for most of the templates tested and has the potential to enable a simple one-pot, single-step co-transcriptional enzymatic capping for mRNA synthesis.

## Methods

### Plasmid construction

The FCE::T7RNAP fusion was constructed by linking the FCE ORF to the 5′ end of T7RNAP ORF via a seven amino acids linker. The FCE::T7RNAP sequence was inserted into a modified pET29a vector at NdeI and SalI sites where the T7 promoter was replaced by a tac promoter. The plasmid was constructed commercially (Genscript).

The 5′UTR panel of IVT transcripts was constructed by inserting the 5′UTR sequence of a synthetic FLuc construct, human hemoglobin beta subunit (HBB, Genbank accession: NM_000518) or pRNA21 5′ to a consensus mammalian Kozak sequence (GCCACC), followed by the start codon of firefly luciferase gene (modified to remove BspQI site), a 3′UTR derived from the publicly available sequence of the BNT162b2 sequence and a poly(A) tail of 60 adenosines and a BspQI site to pUC57 (see Supporting Data). A construct that transcribes a modified form of mRNA-1273 (Moderna) derived from a publicly available sequence ^29^. The sequence was inserted into pUC57 downstream of the consensus mammalian Kozak sequence (above) and upstream of a BsmBI site (see Supporting Data). The plasmids were constructed commercially (Genscript) and linearized by the appropriate Type IIS enzyme (BspQI or BsmBI) and purified by Monarch RNA Cleanup kit (New England Biolabs) prior to IVT reactions.

### Protein expression and purification

The ptac expression construct was used to transform BL21 star DE3. Protein expression was induced by 1 mM IPTG at 37°C when the culture reached an OD600 of 0.6-0.8. The protein was purified by a Ni affinity column followed by size exclusion chromatography using a Superdex 200 column. The protein purity was >90% as judged by Comassie blue-stained SDS-polyacrylamide gel (Supporting data 11). The protein expression and purification were done commercially (Genscript)

### Co-transcriptional capping reactions

IVT and co-transcriptional capping reactions were performed in 30 μL reactions containing 1x T7 RNA polymerase buffer (40 mM Tris-HCl, 20 mM MgCl2, 1 mM DTT, 2 mM spermidine, pH 7.9; New England Biolabs), 5 mM each NTPs, 5 U/mL of E. coli inorganic pyrophosphatase (New England Biolabs), 1 U/mL murine RNase inhibitor (New England Biolabs), 60 nM linearized plasmid template, 0.4 mM SAM, with the indicated concentration of T7 RNA polymerase (New England Biolabs) FCE, VCE or FCE::T7RNAP fusion and 5 U/μL vaccinia cap 2′-O-methyltransferase (New England Biolabs). Reactions were carried out at indicated temperatures for 1 h. An additional 5 mM DTT was added to the reactions to help preserve T7 RNA polymerase activity.

### Post-transcriptional analyses Yield

3 μL of the transcription and co-transcriptional reactions were added to a 6 μL mixture containing 1x DNase I reaction buffer and 0.5 U/μL DNase I (New England Biolabs). After incubation at 37°C for 30 min, the RNA concentration was determined by using the Qubit RNA BR assay.

### RNA cap incorporation

RNA cap incorporation was analyzed by a recently published method^16^. Briefly, for each of the RNA construct, the co-transcriptional reactions were heated at 80°C for 30 seconds with 1 μL of 50 μM of a DNA oligonucleotide designed to anneal to the first 18-22 nt of the specific transcript and contain a desthiobiotin group at the 3′ end. The reactions were allowed to cool down to 25°C at a rate of 0.1°C/s. The reactions were adjusted to 80 μL containing 1x NEBuffer r1.1 (New England Biolabs) and a final concentration of 1.4 nM of hRNase 4 and 0.15 U/μL of polynucleotide kinase (New England Biolabs). The reactions were incubated at 37°C for 1 h and stopped by addition of murine RNase inhibitor to a final concentration of 2 U/μL and incubation at room temperature for 10 minutes. The duplex hRNase 4 cleavage products (5′ fragments of the RNA) were enriched using Hydrophilic Streptavidin Magnetic Beads (New England Biolabs). The beads were washed twice by a high salt buffer (5 mM Tris HCl, pH 7.5, 500 mM NaCl, 0.5 mM EDTA) and thrice using a low salt buffer (5 mM Tris HCl, pH 7.5, 60 mM NaCl, 0.5 mM EDTA). The desthiobiotinylated DNA-5′ cleavage fragment duplexes were released from the streptavidin beads by heating at 65°C for 5 minutes in 35 μL of nuclease-free water. The released RNA-DNA duplexes were further filtered through 0.2 μm Ultrafree-MC Centrifugal Filter (Millipore Sigma) and analyzed by LC-MS/MS ^16^.

