## Supplemental Materials for "Co-transcriptional capping using an RNA capping enzyme-T7 RNA polymerase fusion protein"

### Design of the IVT DNA templates

T7 promoter phi10 (highlighted in gray) was used to drive transcription of RNA initiating with a guanosine (FLuc 5'UTR, pRNA21 5'UTR and mRNA-1273*) whereas T7 phi2.5 promoter (highlighted in gray) was used to drive transcription of HBB 5'UTR, which initiated with an adenosine. The 5'UTR of FLuc, HBB and pRNA21 are shown in bold face. The restriction site that facilitates linearization of the plasmid template (BspQI for FLuc, HBB and pRNA21 5'UTR-FLuc transcripts; BsmBI for mRNA-1273*) are underlined. The T7 transcription cassette was inserted into pUC57 at NdeI and PciI sites (shown in italicized face).

| FLuc 5'UTR-FLuc |
| --- |
| *CATATG*GCTAGCCCCGCGAAATTAATACGACTCACTATA**GGGTCTAGAAATAATTTTGTTTAA**<br>**CTTTAAGAAGGAGATATAACCATGAAAATCGAAGAA**GCCACCATGGAGGACGCCAAGAACATC<br>AAGAAGGGCCCCGCCCCCTTCTACCCCCTGGAGGACGGCACCGCCGGCGAGCAGCTGCACAAG<br>GCCATGAAGCGGTACGCCCTGGTGCCCGGCACCATCGCCTTCACCGACGCCCACATCGAGGTG<br>GACATCACCTACGCCGAGTACTTCGAGATGAGCGTGCGGCTGGCCGAGGCCATGAAGCGGTAC<br>GGCCTGAACACCAACCACCGGATCGTGGTGTGCAGCGAGAACAGCCTGCAGTTCTTCATGCCC<br>GTGCTGGGCGCCCTGTTCATCGGCGTGGCCGTGGCCCCCGCCAACGACATCTACAACGAGCGG<br>GAGCTGCTGAACAGCATGGGCATCAGCCAGCCCACCGTGGTGTTCGTGAGCAAGAAGGGCCTG<br>CAGAAGATCCTGAACGTGCAGAAGAAGCTGCCCATCATCCAGAAGATCATCATCATGGACAGC<br>AAGACCGACTACCAGGGCTTCCAGAGCATGTACACCTTCGTGACCAGCCACCTGCCCCCCGGC<br>TTCAACGAGTACGACTTCGTGCCCGAGAGCTTCGACCGGGACAAGACCATCGCCCTGATCATG<br>AACAGCAGCGGCAGCACCGGCCTGCCCAAGGGCGTGGCCCTGCCCCACCGGACCGCCTGCGTG<br>CGGTTCAGCCACGCCCGGGACCCCATCTTCGGCAACCAGATCATCCCCGACACCGCCATCCTG<br>AGCGTGGTGCCCTTCCACCACGGCTTCGGCATGTTCACCACCCTGGGCTACCTGATCTGCGGC<br>TTCCGGGTGGTGCTGATGTACCGGTTCGAGGAGGAGCTGTTCCTGCGGAGCCTGCAGGACTAC |

AAGATCCAGAGCGCCCTGCTGGTGCCCACCCTGTTCAGCTTCTTCGCCAAGAGCACCCTGATC
GACAAGTACGACCTGAGCAACCTGCACGAGATCGCCAGCGGCGGCGCCCCCCTGAGCAAGGAG
GTGGGCGAGGCCGTGGCCAAGCGGTTCCACCTGCCCGGCATCCGGCAGGGCTACGGCCTGACC
GAGACCACCAGCGCCATCCTGATCACCCCCGAGGGCGACGACAAGCCCGGCGCCGTGGGCAAG
GTGGTGCCCTTCTTCGAGGCCAAGGTGGTGGACCTGGACACCGGCAAGACCCTGGGCGTGAAC
CAGCGGGGCGAGCTGTGCGTGCGGGGCCCCATGATCATGAGCGGCTACGTGAACAACCCCGAG
GCCACCAACGCCCTGATCGACAAGGACGGCTGGCTGCACAGCGGCGACATCGCCTACTGGGAC
GAGGACGAGCACTTCTTCATCGTGGACCGGCTGAAAAGTCTGATCAAGTACAAGGGCTACCAG
GTGGCCCCCGCCGAGCTGGAGAGCATCCTGCTGCAGCACCCCAACATCTTCGACGCCGGCGTG
GCCGGCCTGCCCGACGACGACGCCGGCGAGCTGCCCGCCGCCGTGGTGGTGCTGGAGCACGGC
AAGACCATGACCGAGAAGGAGATCGTGGACTACGTGGCCAGCCAGGTGACCACCGCCAAGAAG
CTGCGGGGCGGCGTGGTGTTCGTGGACGAGGTGCCCAAGGGCCTGACCGGCAAGCTGGACGCC
CGGAAGATCCGGGAGATCCTGATCAAGGCCAAGAAGGGCGGCAAGATCGCCGTGTGAGGCGCG
CCTATGTTACGTGCAAAGGTGATTGTCACCCCCCGAAAGACCATATTGTGACACACCCTCAGT
ATCACGCCCAAACATTTACAGCCGCGGTGTCAAAAACCGCGTGGACGTGGTTAACATCCCTGC
TGGGAGGATCAGCCGTAATTATTATAATTGGCTTGGTGCTGGCTACTATTGTGGCCATGTACG
TGCTGACCAACCAGAAACATAATTGAATACAGCAGCAATTGGCAAGCTGCTTACATAGAACTC
GCGGCGATTGGCATGCCGCCTTAAAATTTTTATTTTATTTTTCTTTTCTTTTCCGAATCGGAT
TTTGTTTTTAATATTTCAAAAAAAAAAAAAAAAAAAAAAAAAAAAAAAAAAAAAAAAAAAAAA
AAAAAAAAAAAAAAAGAAGAGC*ACATGT*

HBB 5’UTR-FLuc

*CATATG*TAATACGACTCACTATT**ACATTTGCTTCTGACACAACTGTGTTCACTAGCAACCTCA**
**AACAGACACC**GCCACCATGGAGGACGCCAAGAACATCAAGAAGGGCCCCGCCCCCTTCTACCC
CCTGGAGGACGGCACCGCCGGCGAGCAGCTGCACAAGGCCATGAAGCGGTACGCCCTGGTGCC
CGGCACCATCGCCTTCACCGACGCCCACATCGAGGTGGACATCACCTACGCCGAGTACTTCGA
GATGAGCGTGCGGCTGGCCGAGGCCATGAAGCGGTACGGCCTGAACACCAACCACCGGATCGT
GGTGTGCAGCGAGAACAGCCTGCAGTTCTTCATGCCCGTGCTGGGCGCCCTGTTCATCGGCGT
GGCCGTGGCCCCCGCCAACGACATCTACAACGAGCGGGAGCTGCTGAACAGCATGGGCATCAG
CCAGCCCACCGTGGTGTTCGTGAGCAAGAAGGGCCTGCAGAAGATCCTGAACGTGCAGAAGAA
GCTGCCCATCATCCAGAAGATCATCATCATGGACAGCAAGACCGACTACCAGGGCTTCCAGAG

CATGTACACCTTCGTGACCAGCCACCTGCCCCCCGGCTTCAACGAGTACGACTTCGTGCCCGA
GAGCTTCGACCGGGACAAGACCATCGCCCTGATCATGAACAGCAGCGGCAGCACCGGCCTGCC
CAAGGGCGTGGCCCTGCCCCACCGGACCGCCTGCGTGCGGTTCAGCCACGCCCGGGACCCCAT
CTTCGGCAACCAGATCATCCCCGACACCGCCATCCTGAGCGTGGTGCCCTTCCACCACGGCTT
CGGCATGTTCACCACCCTGGGCTACCTGATCTGCGGCTTCCGGGTGGTGCTGATGTACCGGTT
CGAGGAGGAGCTGTTCCTGCGGAGCCTGCAGGACTACAAGATCCAGAGCGCCCTGCTGGTGCC
CACCCTGTTCAGCTTCTTCGCCAAGAGCACCCTGATCGACAAGTACGACCTGAGCAACCTGCA
CGAGATCGCCAGCGGCGGCGCCCCCCTGAGCAAGGAGGTGGGCGAGGCCGTGGCCAAGCGGTT
CCACCTGCCCGGCATCCGGCAGGGCTACGGCCTGACCGAGACCACCAGCGCCATCCTGATCAC
CCCCGAGGGCGACGACAAGCCCGGCGCCGTGGGCAAGGTGGTGCCCTTCTTCGAGGCCAAGGT
GGTGGACCTGGACACCGGCAAGACCCTGGGCGTGAACCAGCGGGGCGAGCTGTGCGTGCGGGG
CCCCATGATCATGAGCGGCTACGTGAACAACCCCGAGGCCACCAACGCCCTGATCGACAAGGA
CGGCTGGCTGCACAGCGGCGACATCGCCTACTGGGACGAGGACGAGCACTTCTTCATCGTGGA
CCGGCTGAAAAGTCTGATCAAGTACAAGGGCTACCAGGTGGCCCCCGCCGAGCTGGAGAGCAT
CCTGCTGCAGCACCCCAACATCTTCGACGCCGGCGTGGCCGGCCTGCCCGACGACGACGCCGG
CGAGCTGCCCGCCGCCGTGGTGGTGCTGGAGCACGGCAAGACCATGACCGAGAAGGAGATCGT
GGACTACGTGGCCAGCCAGGTGACCACCGCCAAGAAGCTGCGGGGCGGCGTGGTGTTCGTGGA
CGAGGTGCCCAAGGGCCTGACCGGCAAGCTGGACGCCCGGAAGATCCGGGAGATCCTGATCAA
GGCCAAGAAGGGCGGCAAGATCGCCGTGTGAGGCGCGCCTATGTTACGTGCAAAGGTGATTGT
CACCCCCCGAAAGACCATATTGTGACACACCCTCAGTATCACGCCCAAACATTTACAGCCGCG
GTGTCAAAAACCGCGTGGACGTGGTTAACATCCCTGCTGGGAGGATCAGCCGTAATTATTATA
ATTGGCTTGGTGCTGGCTACTATTGTGGCCATGTACGTGCTGACCAACCAGAAACATAATTGA
ATACAGCAGCAATTGGCAAGCTGCTTACATAGAACTCGCGGCGATTGGCATGCCGCCTTAAAA
TTTTTATTTTATTTTTCTTTTCTTTTCCGAATCGGATTTTGTTTTTAATATTTCAAAAAAAAA
AAAAAAAAAAAAAAAAAAAAAAAAAAAAAAAAAAAAAAAAAAAAAAAAAAAAGAAGAGC*ACAT*
*GT*

pRNA21 5'UTR-FLuc

*CATATG*GCTAGCCCCGCGAAATTAATACGACTCACTATA**GGGAAATAAGAGAGAAAAGAAGAG**
**TAAGAAGAAATATAAGAGCCACC**GCCACCATGGAGGACGCCAAGAACATCAAGAAGGGCCCCG
CCCCCTTCTACCCCCTGGAGGACGGCACCGCCGGCGAGCAGCTGCACAAGGCCATGAAGCGGT

ACGCCCTGGTGCCCGGCACCATCGCCTTCACCGACGCCCACATCGAGGTGGACATCACCTACG
CCGAGTACTTCGAGATGAGCGTGCGGCTGGCCGAGGCCATGAAGCGGTACGGCCTGAACACCA
ACCACCGGATCGTGGTGTGCAGCGAGAACAGCCTGCAGTTCTTCATGCCCGTGCTGGGCGCCC
TGTTCATCGGCGTGGCCGTGGCCCCCGCCAACGACATCTACAACGAGCGGGAGCTGCTGAACA
GCATGGGCATCAGCCAGCCCACCGTGGTGTTCGTGAGCAAGAAGGGCCTGCAGAAGATCCTGA
ACGTGCAGAAGAAGCTGCCCATCATCCAGAAGATCATCATCATGGACAGCAAGACCGACTACC
AGGGCTTCCAGAGCATGTACACCTTCGTGACCAGCCACCTGCCCCCCGGCTTCAACGAGTACG
ACTTCGTGCCCGAGAGCTTCGACCGGGACAAGACCATCGCCCTGATCATGAACAGCAGCGGCA
GCACCGGCCTGCCCAAGGGCGTGGCCCTGCCCCACCGGACCGCCTGCGTGCGGTTCAGCCACG
CCCGGGACCCCATCTTCGGCAACCAGATCATCCCCGACACCGCCATCCTGAGCGTGGTGCCCT
TCCACCACGGCTTCGGCATGTTCACCACCCTGGGCTACCTGATCTGCGGCTTCCGGGTGGTGC
TGATGTACCGGTTCGAGGAGGAGCTGTTCCTGCGGAGCCTGCAGGACTACAAGATCCAGAGCG
CCCTGCTGGTGCCCACCCTGTTCAGCTTCTTCGCCAAGAGCACCCTGATCGACAAGTACGACC
TGAGCAACCTGCACGAGATCGCCAGCGGCGGCGCCCCCCTGAGCAAGGAGGTGGGCGAGGCCG
TGGCCAAGCGGTTCCACCTGCCCGGCATCCGGCAGGGCTACGGCCTGACCGAGACCACCAGCG
CCATCCTGATCACCCCCGAGGGCGACGACAAGCCCGGCGCCGTGGGCAAGGTGGTGCCCTTCT
TCGAGGCCAAGGTGGTGGACCTGGACACCGGCAAGACCCTGGGCGTGAACCAGCGGGGCGAGC
TGTGCGTGCGGGGCCCCATGATCATGAGCGGCTACGTGAACAACCCCGAGGCCACCAACGCCC
TGATCGACAAGGACGGCTGGCTGCACAGCGGCGACATCGCCTACTGGGACGAGGACGAGCACT
TCTTCATCGTGGACCGGCTGAAAAGTCTGATCAAGTACAAGGGCTACCAGGTGGCCCCCGCCG
AGCTGGAGAGCATCCTGCTGCAGCACCCCAACATCTTCGACGCCGGCGTGGCCGGCCTGCCCG
ACGACGACGCCGGCGAGCTGCCCGCCGCCGTGGTGGTGCTGGAGCACGGCAAGACCATGACCG
AGAAGGAGATCGTGGACTACGTGGCCAGCCAGGTGACCACCGCCAAGAAGCTGCGGGGCGGCG
TGGTGTTCGTGGACGAGGTGCCCAAGGGCCTGACCGGCAAGCTGGACGCCCGGAAGATCCGGG
AGATCCTGATCAAGGCCAAGAAGGGCGGCAAGATCGCCGTGTGAGGCGCGCCTATGTTACGTG
CAAAGGTGATTGTCACCCCCCGAAAGACCATATTGTGACACACCCTCAGTATCACGCCCAAAC
ATTTACAGCCGCGGTGTCAAAAACCGCGTGGACGTGGTTAACATCCCTGCTGGGAGGATCAGC
CGTAATTATTATAATTGGCTTGGTGCTGGCTACTATTGTGGCCATGTACGTGCTGACCAACCA
GAAACATAATTGAATACAGCAGCAATTGGCAAGCTGCTTACATAGAACTCGCGGCGATTGGCA
TGCCGCCTTAAAATTTTTATTTTATTTTTCTTTTCTTTTCCGAATCGGATTTTGTTTTTAATA

| |
|---|
| TTTCAAAAAAAAAAAAAAAAAAAAAAAAAAAAAAAAAAAAAAAAAAAAAAAAAAAAAAAA<br>AAGAAGAGC*ACATGT* |

| mRNA-1273* |
|---|
| *CATATG*GCTAGCCCCGCGAAATTAATACGACTCACTATAGGGAAATAAGAGAGAAAAGAAGAG<br>TAAGAAGAAATATAAGACCCCGGCGCCGCCACCATGTTCGTGTTCCTGGTGCTGCTGCCCCTG<br>GTGAGCAGCCAGTGCGTGAACCTGACCACCCGGACCCAGCTGCCACCAGCCTACACCAACAGC<br>TTCACCCGGGGCGTCTACTACCCCGACAAGGTGTTCCGGAGCAGCGTCCTGCACAGCACCCAG<br>GACCTGTTCCTGCCCTTCTTCAGCAACGTGACCTGGTTCCACGCCATCCACGTGAGCGGCACC<br>AACGGCACCAAGCGGTTCGACAACCCCGTGCTGCCCTTCAACGACGGCGTGTACTTCGCCAGC<br>ACCGAGAAGAGCAACATCATCCGGGGCTGGATCTTCGGCACCACCCTGGACAGCAAGACCCAG<br>AGCCTGCTGATCGTGAATAACGCCACCAACGTGGTGATCAAGGTGTGCGAGTTCCAGTTCTGC<br>AACGACCCCTTCCTGGGCGTGTACTACCACAAGAACAACAAGAGCTGGATGGAGAGCGAGTTC<br>CGGGTGTACAGCAGCGCCAACAACTGCACCTTCGAGTACGTGAGCCAGCCCTTCCTGATGGAC<br>CTGGAGGGCAAGCAGGGCAACTTCAAGAACCTGCGGGAGTTCGTGTTCAAGAACATCGACGGC<br>TACTTCAAGATCTACAGCAAGCACACCCCAATCAACCTGGTGCGGGATCTGCCCCAGGGCTTC<br>TCAGCCCTGGAGCCCCTGGTGGACCTGCCCATCGGCATCAACATCACCCGGTTCCAGACCCTG<br>CTGGCCCTGCACCGGAGCTACCTGACCCCAGGCGACAGCAGCAGCGGGTGGACAGCAGGCGCG<br>GCTGCTTACTACGTGGGCTACCTGCAGCCCCGGACCTTCCTGCTGAAGTACAACGAGAACGGC<br>ACCATCACCGACGCCGTGGACTGCGCCCTGGACCCTCTGAGCGAGACCAAGTGCACCCTGAAG<br>AGCTTCACCGTGGAGAAGGGCATCTACCAGACCAGCAACTTCCGGGTGCAGCCCACCGAGAGC<br>ATCGTGCGGTTCCCCAACATCACCAACCTGTGCCCCTTCGGCGAGGTGTTCAACGCCACCCGG<br>TTCGCCAGCGTGTACGCCTGGAACCGGAAGCGGATCAGCAACTGCGTGGCCGACTACAGCGTG<br>CTGTACAACAGCGCCAGCTTCAGCACCTTCAAGTGCTACGGCGTGAGCCCCACCAAGCTGAAC<br>GACCTGTGCTTCACCAACGTGTACGCCGACAGCTTCGTGATCCGTGGCGACGAGGTGCGGCAG<br>ATCGCACCCGGCCAGACAGGCAAGATCGCCGACTACAACTACAAGCTGCCCGACGACTTCACC<br>GGCTGCGTGATCGCCTGGAACAGCAACAACCTCGACAGCAAGGTGGGCGGCAACTACAACTAC<br>CTGTACCGGCTGTTCCGGAAGAGCAACCTGAAGCCCTTCGAGCGGGACATCAGCACCGAGATC<br>TACCAAGCCGGCTCCACCCCTTGCAACGGCGTGGAGGGCTTCAACTGCTACTTCCCTCTGCAG<br>AGCTACGGCTTCCAGCCCACCAACGGCGTGGGCTACCAGCCCTACCGGGTGGTGGTGCTGAGC<br>TTCGAGCTGCTGCACGCCCCAGCCACCGTGTGTGGCCCCAAGAAGAGCACCAACCTGGTGAAG |

AACAAGTGCGTGAACTTCAACTTCAACGGCCTTACCGGCACCGGCGTGCTGACCGAGAGCAAC
AAGAAATTCCTGCCCTTTCAGCAGTTCGGCCGGGACATCGCCGACACCACCGACGCTGTGCGG
GATCCCCAGACCCTGGAGATCCTGGACATCACCCCTTGCAGCTTCGGCGGCGTGAGCGTGATC
ACCCCAGGCACCAACACCAGCAACCAGGTGGCCGTGCTGTACCAGGACGTGAACTGCACCGAG
GTGCCCGTGGCCATCCACGCCGACCAGCTGACACCCACCTGGCGGGTCTACAGCACCGGCAGC
AACGTGTTCCAGACCCGGGCCGGTTGCCTGATCGGCGCCGAGCACGTGAACAACAGCTACGAG
TGCGACATCCCCATCGGCGCCGGCATCTGTGCCAGCTACCAGACCCAGACCAATTCACCCCGG
AGGGCAAGGAGCGTGGCCAGCCAGAGCATCATCGCCTACACCATGAGCCTGGGCGCCGAGAAC
AGCGTGGCCTACAGCAACAACAGCATCGCCATCCCCACCAACTTCACCATCAGCGTGACCACC
GAGATTCTGCCCGTGAGCATGACCAAGACCAGCGTGGACTGCACCATGTACATCTGCGGCGAC
AGCACCGAGTGCAGCAACCTGCTGCTGCAGTACGGCAGCTTCTGCACCCAGCTGAACCGGGCC
CTGACCGGCATCGCCGTGGAGCAGGACAAGAACACCCAGGAGGTGTTCGCCCAGGTGAAGCAG
ATCTACAAGACCCCTCCCATCAAGGACTTCGGCGGCTTCAACTTCAGCCAGATCCTGCCCGAC
CCCAGCAAGCCCAGCAAGCGGAGCTTCATCGAGGACCTGCTGTTCAACAAGGTGACCCTAGCC
GACGCCGGCTTCATCAAGCAGTACGGCGACTGCCTCGGCGACATAGCCGCCCGGGACCTGATC
TGCGCCCAGAAGTTCAACGGCCTGACCGTGCTGCCTCCCCTGCTGACCGACGAGATGATCGCC
CAGTACACCAGCGCCCTGTTAGCCGGAACCATCACCAGCGGCTGGACTTTCGGCGCTGGAGCC
GCTCTGCAGATCCCCTTCGCCATGCAGATGGCCTACCGGTTCAACGGCATCGGCGTGACCCAG
AACGTGCTGTACGAGAACCAGAAGCTGATCGCCAACCAGTTCAACAGCGCCATCGGCAAGATC
CAGGACAGCCTGAGCAGCACCGCTAGCGCCCTGGGCAAGCTGCAGGACGTGGTGAACCAGAAC
GCCCAGGCCCTGAACACCCTGGTGAAGCAGCTGAGCAGCAACTTCGGCGCCATCAGCAGCGTG
CTGAACGACATCCTGAGCCGGCTGGACCCTCCCGAGGCCGAGGTGCAGATCGACCGGCTGATC
ACTGGCCGGCTGCAGAGCCTGCAGACCTACGTGACCCAGCAGCTGATCCGGGCCGCCGAGATT
CGGGCCAGCGCCAACCTGGCCGCCACCAAGATGAGCGAGTGCGTGCTGGGCCAGAGCAAGCGG
GTGGACTTCTGCGGCAAGGGCTACCACCTGATGAGCTTTCCCCAGAGCGCACCCCACGGAGTG
GTGTTCCTGCACGTGACCTACGTGCCCGCCCAGGAGAAGAACTTCACCACCGCCCCAGCCATC
TGCCACGACGGCAAGGCCCACTTTCCCCGGGAGGGCGTGTTCGTGAGCAACGGCACCCACTGG
TTCGTGACCCAGCGGAACTTCTACGAGCCCCAGATCATCACCACCGACAACACCTTCGTGAGC
GGCAACTGCGACGTGGTGATCGGCATCGTGAACAACACCGTGTACGATCCCCTGCAGCCCGAG
CTGGACAGCTTCAAGGAGGAGCTGGACAAGTACTTCAAGAATCACACCAGCCCCGACGTGGAC
CTGGGCGACATCAGCGGCATCAACGCCAGCGTGGTGAACATCCAGAAGGAGATCGATCGGCTG

```
AACGAGGTGGCCAAGAACCTGAACGAGAGCCTGATCGACCTGCAGGAGCTGGGCAAGTACGAG
CAGTACATCAAGTGGCCCTGGTACATCTGGCTGGGCTTCATCGCCGGCCTGATCGCCATCGTG
ATGGTGACCATCATGCTGTGCTGCATGACCAGCTGCTGCAGCTGCCTGAAGGGCTGTTGCAGC
TGCGGCAGCTGCTGCAAGTTCGACGAGGACGACAGCGAGCCCGTGCTGAAGGGCGTGAAGCTG
CACTACACCTGATAATAGGCTGGAGCCTCGGTGGCCTAGCTTCTTGCCCCTTGGGCCTCCCCC
CAGCCCCTCCTCCCCTTCCTGCACCCGTACCCCCGTGGTCTTTGAATAAAGTCTGAGTGGCG
GCAAAAAAAAAAAAAAAAAAAAAAAAAAAAAAAAAAAAAAAAAAAAAAAAAAAAAAAAAA
AAAAAAAAAAAAAAAAAAAAAAAAAAAAAAAAAAAAAAGAGACGACATGT
```

**Supporting Data 1**

Co-transcriptional capping using separate T7 RNA polymerase and RNA capping enzymes. Under standard IVT conditions, 20 µL reactions containing 1x T7 RNA polymerase buffer (40 mM Tris-HCl, 20 mM MgCl2, 1 mM DTT, 2 mM spermidine, pH 7.9; New England Biolabs), 5 mM each NTPs, 5 U/mL of E. coli inorganic pyrophosphatase (New England Biolabs), 1 U/mL murine RNase inhibitor (New England Biolabs), 80 nM T7 RNA polymerase (New England Biolabs), 120 nM linearized PCR-amplified FLuc IVT template DNA containing the T7 pih10 promoter, 0.4 mM SAM and indicated concentrations of VCE or FCE were incubated at 37°C or 45°C for 1 h. The yield of transcription was measured using the method described in Methods. The level of cap incorporation was measured using a RNase H-based LC-MS/MS method[1]. It required up to 250 nM of VCE or 500 nM of FCE at 45°C to achieve ~95% Cap-0 incorporation in the in vitro transcripts. Reactions performed at 37°C resulted in <60% Cap-0 incorporation (data not shown).

**Supporting Data 2**

RNA capping activity of the FCE::T7RNAP fusion protein. Indicated concentrations of FCE or the FCE::T7RNAP fusion protein was incubated in 10 µL reactions containing 1x FCE RNA capping buffer (50 mM Tris-HCl, pH 8.0, 5 mM KCl,1 mM $MgCl_2$, 1 mM DTT, 0.02% Poloxamer 188), 0.1 mM SAM, 0.5 mM GTP, 0.5 µM of a 5' triphosphate 3' FAM-labeled 25 nt synthetic RNA at 37°C for 30 min. After quenching in a final concentration of 0.1% SDS, 10 mM EDTA, the reactions were analyzed by capillary electrophoresis[1]. The results showed that the FCE::T7RNAP fusion exhibited RNA capping activity level highly comparable to FCE.

**Supporting Data 3**

Full dataset of Figure 1b and 1c, showing the percentage of all the intermediate products of the enzymatic capping reactions using a mixture of 100 mM FCE and 80 mM T7RNAP, 100 mM of FCE::T7RNAP or T7RNAP only at 30°C, 37°C and 45°C.

**Supporting Data 4**

FCE::T7RNAP fusion in Cap-1 co-transcriptional capping using PCR-amplified DNA as template. 20 µL reactions containing 1x T7 RNA polymerase buffer (40 mM Tris-HCl, 20 mM MgCl2, 1 mM DTT, 2 mM spermidine, pH 7.9; New England Biolabs), 5 mM each NTPs, 5 U/mL of *E. coli* inorganic pyrophosphatase (New England Biolabs), 1 U/mL murine RNase inhibitor (New England Biolabs), 80 nM T7 RNA polymerase (New England Biolabs) or indicated concentration of the FCE::T7RNAP fusion and 5 U/µL vaccinia cap 2'-O-methyltransferae (New England Biolabs), 120 nM PCR-amplified FLuc IVT template DNA containing the T7 pih10 promoter, and 0.4 mM SAM were incubated at 37°C or 45°C for 1 h. The yield of transcription was measured using the method described in Methods. The level of cap incorporation was measured using a RNase H-based LC-MS/MS method[1]. At 45°C, the FCE::T7RNAP fusion, in conjunction with vaccinia cap 2'-O-methyltransferase, generated 100% Cap-1 RNA at all FCE::T7RNAP concentrations tested. Transcription yield increased as FCE::T7RNAP concentration increased. At 37°C, a maximum of 60% of Cap-1 incorporation was achieved with transcriptional yield lower than reactions containing T7 RNAP. Datapoints are the average values of two independent experiments. Error bars represent one standard deviation.

**Supporting Data 5**

The effect of FCE::T7RNAP concentration and reaction temperature on co-transcriptional capping. Full dataset of Figure 1d showing the percentage of all the intermediate products of the enzymatic capping reactions.

**Supporting Data 6**

Post-transcriptional capping of the 5'UTR panel using FCE. IVT was carried out using 60 nM of linearized plasmid templates and standard IVT conditions (Methods) at 37°C for 1 h with the omission of SAM. The in vitro transcripts were purified using Monarch RNA Clean-up kit (50 µg scale). 0.5 µM of the purified transcripts were incubated with indicated concentrations of FCE in a 20 µL reaction containing 1x FCE capping buffer (50 mM Tris-HCl, pH 8.0, 5 mM KCl,1 mM $MgCl_2$, 1 mM DTT, 0.02% Poloxamer 188), 0.1 mM SAM, and 0.5 mM GTP. After incubation at 37°C for 30 min, the level of Cap-0 incorporation was assessed using the RNase H-based LC-MS/MS method[1]. Transcripts of all three 5'UTR were capped to 94-100% using as low as 50 nM FCE. Datapoints are averaged values of three independent experiments. Error bars represent one standard deviation.

**Supporting Data 7**

Dependence of transcription yield and template concentration. A 94-bp template DNA encoding a 77 nt RNA was synthesized and PCR-amplified. Using the HiScribe T7 Quick High Yield RNA Synthesis Kit (New England Biolabs), increasing concentration of template DNA from 125 nM (7.3 ng/µL) to 4 µM (235 ng/µL) resulted in an exponential increase of yield in a 2-h or overnight time period.

**Supporting Data 8**

FCE::T7RNAP fusion and vaccinia cap 2'-O-methyltransferase in Cap-1 co-transcriptional capping of the 5'UTR panel. Full dataset of Figure 2a, showing the percentage of all the intermediate products of the enzymatic capping reactions.

**Supporting Data 9**

FCE::T7RNAP fusion in Cap-1 co-transcriptional capping of mRNA-1273*. Full dataset of Figure 2b, showing the percentage of all the intermediate products of the enzymatic capping reactions.

**Supporting Data 10**

Time course of co-transcriptional capping using FCE::T7RNAP fusion. (a) The yield of transcription over time for 80 nM T7 RNAP only, 100 nM FCE::T7RNAP fusion protein and a mix containing 100 nM FCE and 80 nM T7 RNAP (FCE + T7NRAP). The reactions were carried out using 60 nM of linearized pRNA21-FLuc template for indicated time periods. RNA quantitation was done using Qubit RNA BR kit and a Qubit spectrophotometer. The numbers are average values of quadrupled experiments with error bar representing one standard deviation. (b) Representative gel images of the IVT products from the time course study. The IVT products were treated with DNase I (see Methods) and analyzed on TapeStation (Agilent) using the RNA ScreenTapes and Reagents according to manufacturer's instructions.

**Supporting Data 11**

SDS-PAGE analysis of the purified FCE::T7RNAP fusion protein. Lane R contains the FCE::T7RNAP fusion purified as described in Method. The purification and SDS-PAGE analysis were performed commercially (Genscript).

**Supporting Data 12**

Gel electrophoresis of IVT (T7 RNAP) or co-transcriptional capping products by FCE::T7RNAP after purification using Monarch RNA Clean-up kit (50 µg scale; New England Biolabs). The RNA FLuc, HBB, pRNA21 5'UTR-FLuc (2 kb) and mRNA-1273* (4.1 kb) were analyzed on 2% E-gel (ThermoFisher). The bands were visualized using the Amersham Typhoon RGB scanner with the Cy2 channel.
