## Supporting Data for "Co-transcriptional capping using an RNA capping enzyme-T7 RNA polymerase fusion protein"

Supporting Data 1

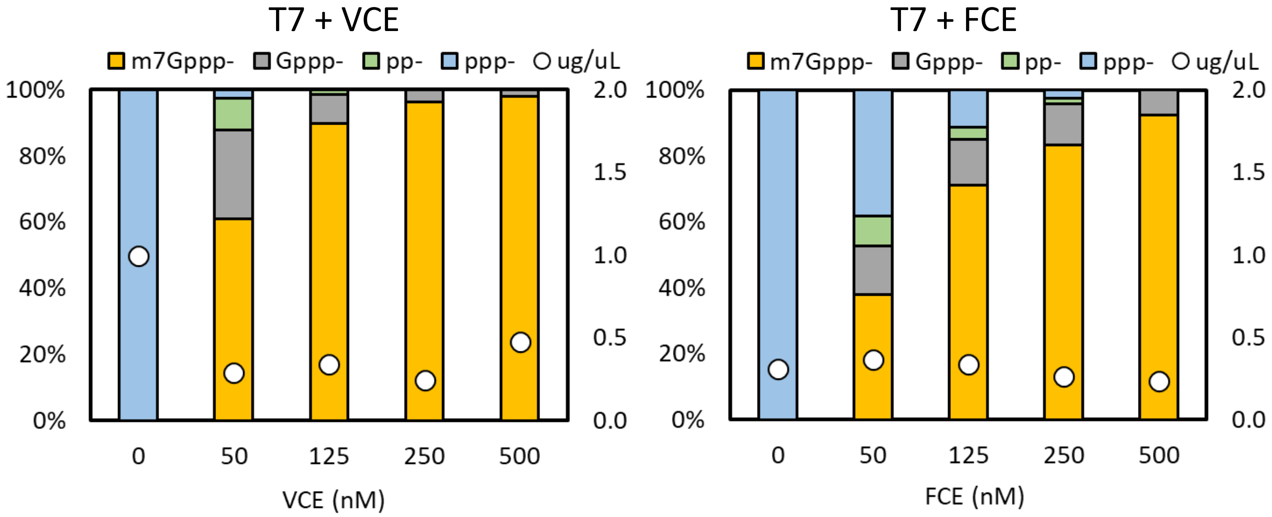

#### Supporting Data 2

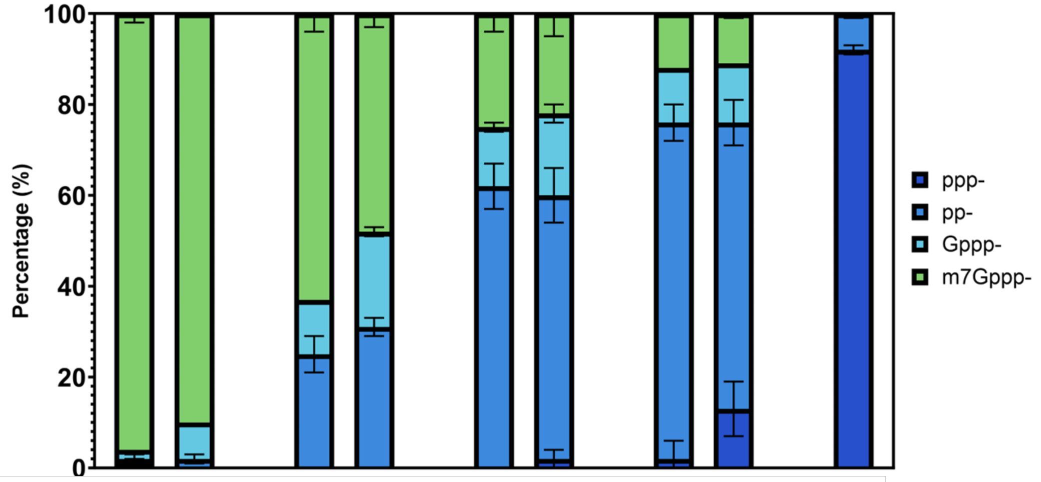


| | | | | | | | | | |
|---|---|---|---|---|---|---|---|---|---|
| FCE | + | - | + | - | + | - | + | - | - |
| FCE::T7RNAP | - | + | - | + | - | + | - | + | - |
| conc. (nM) | 0.625 | | 0.312 | | 0.156 | | 0.078 | | 0 |

### Supporting Data 3

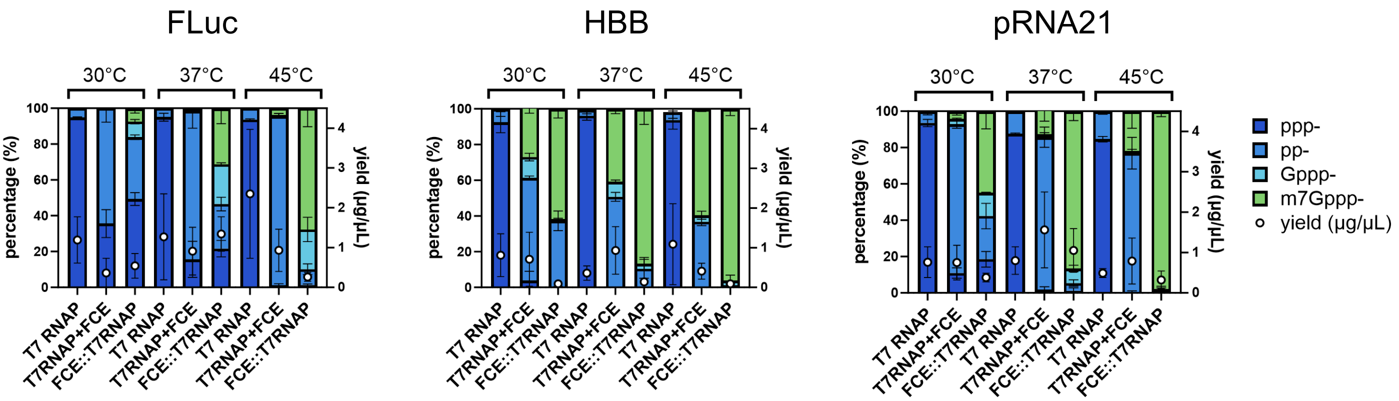

Supporting Data 4

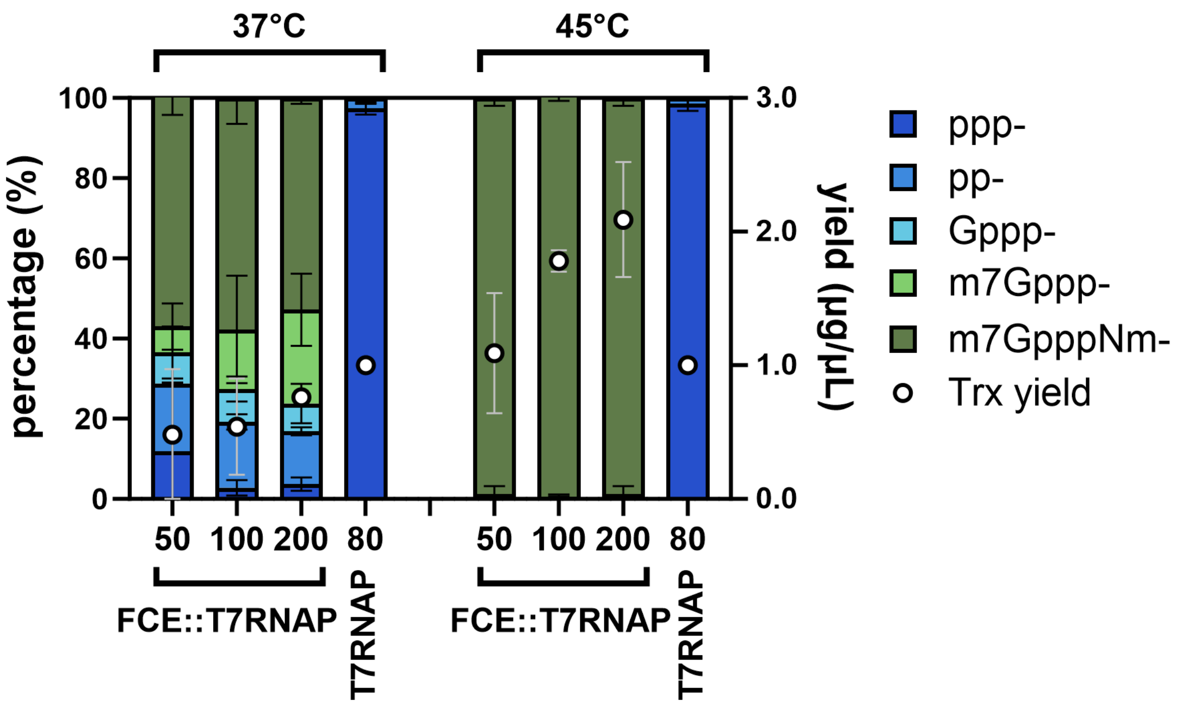

Supporting Data 5

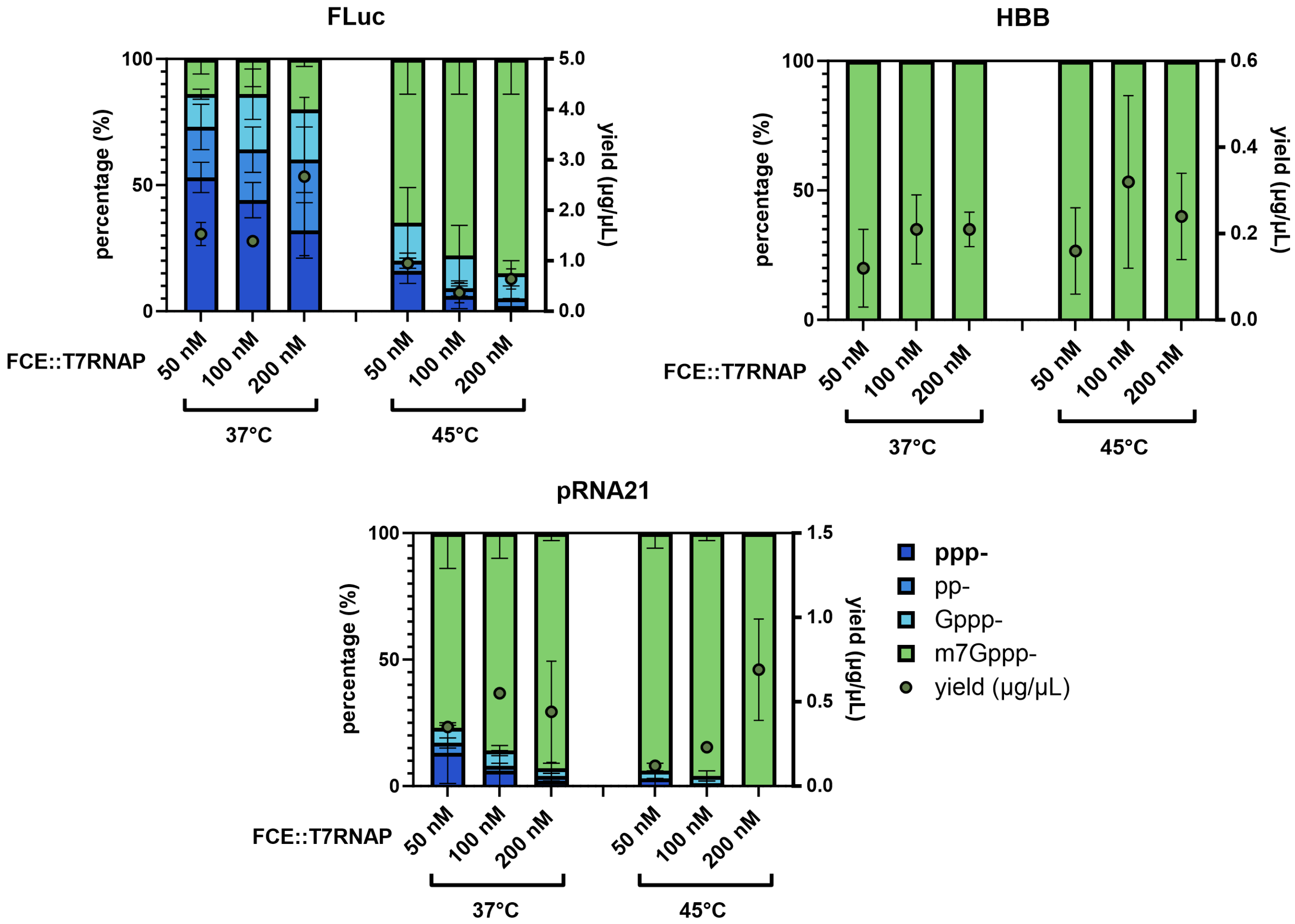

Supporting Data 6

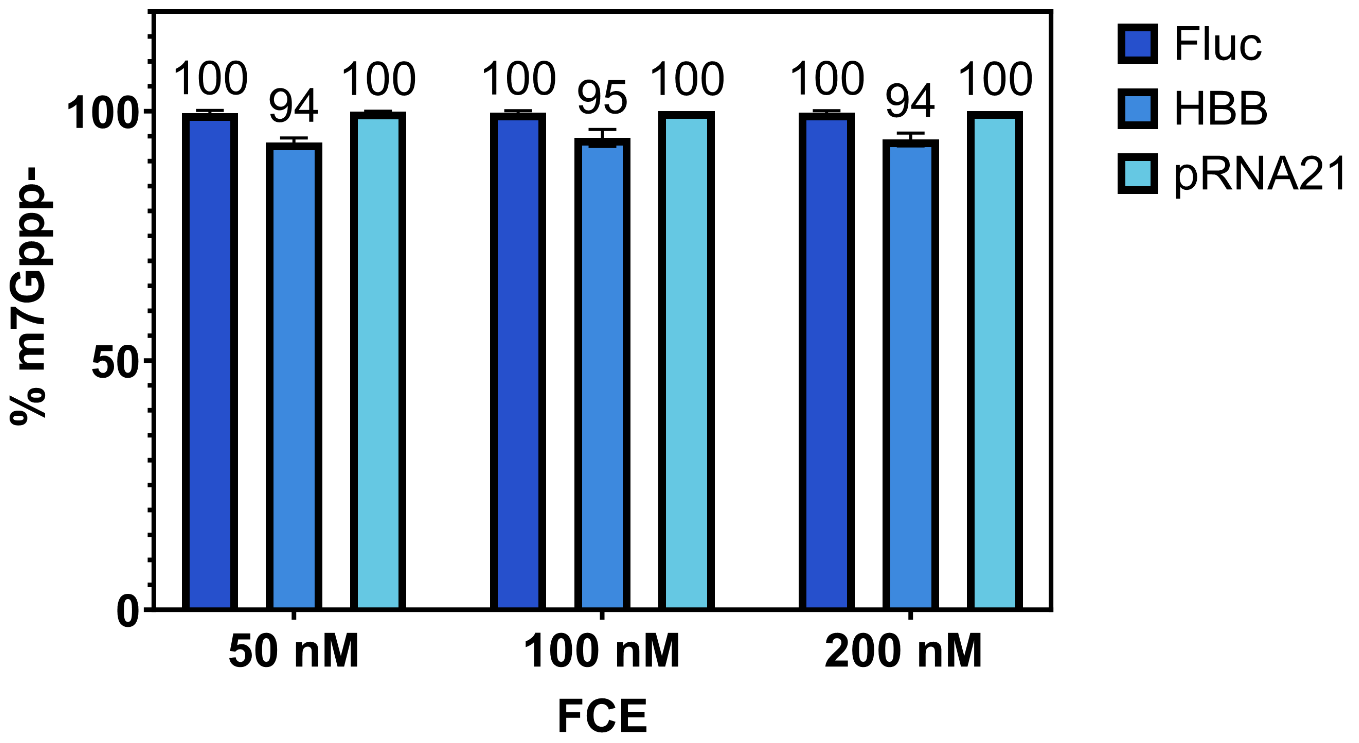

Supporting Data 7

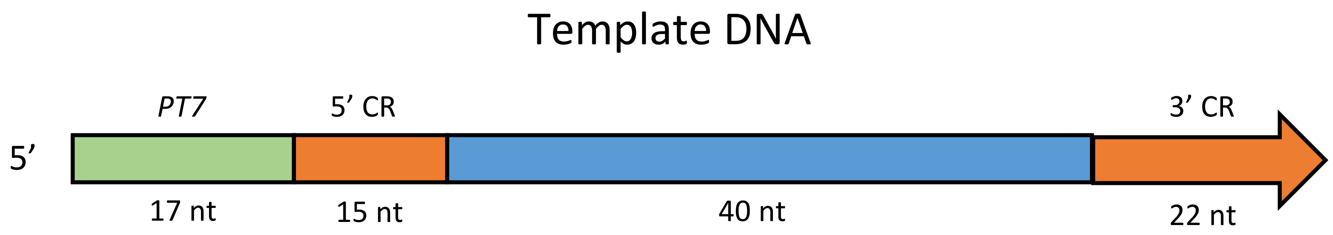


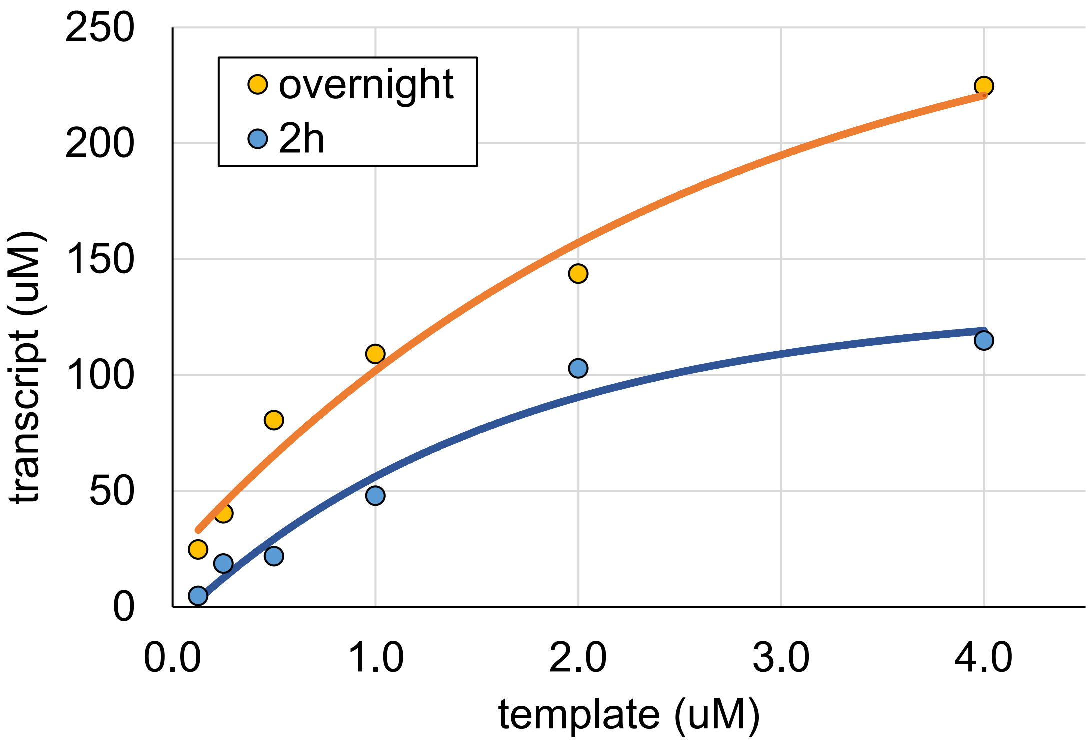

### Supporting Data 8

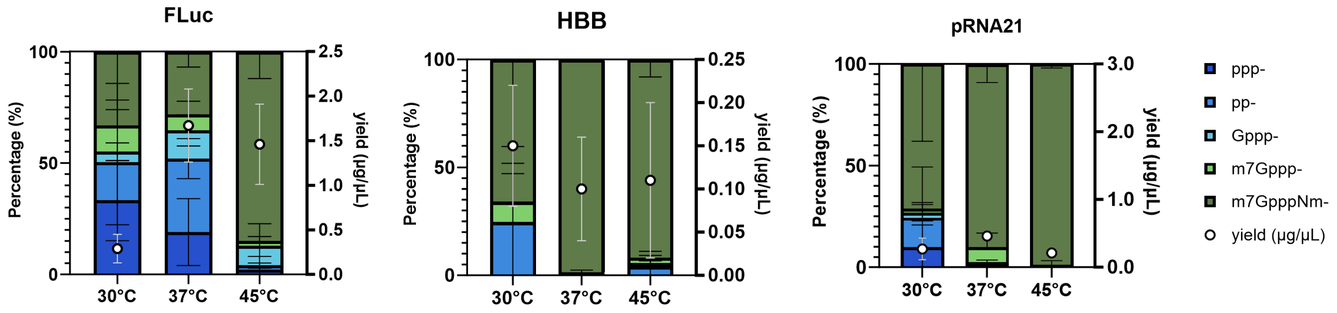

Supporting Data 9

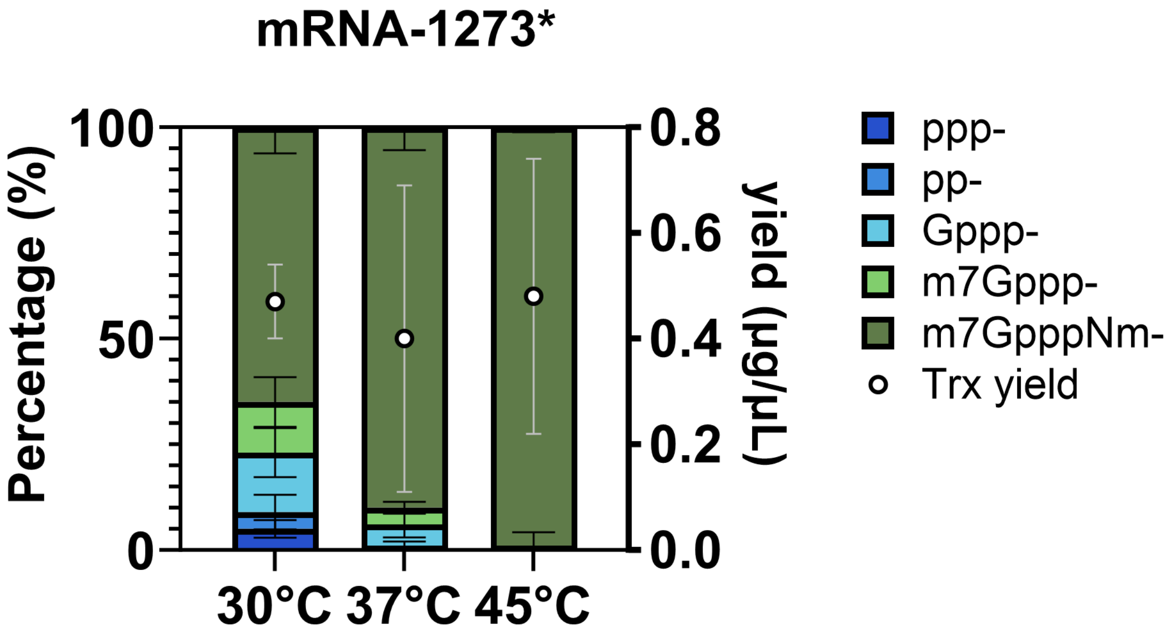

Supporting Data 10

a

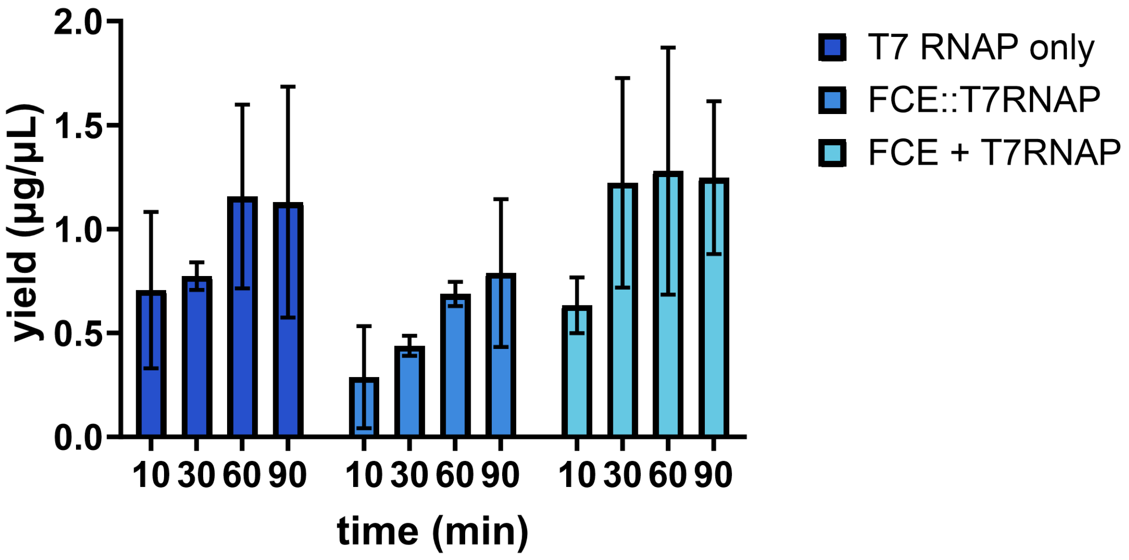


b

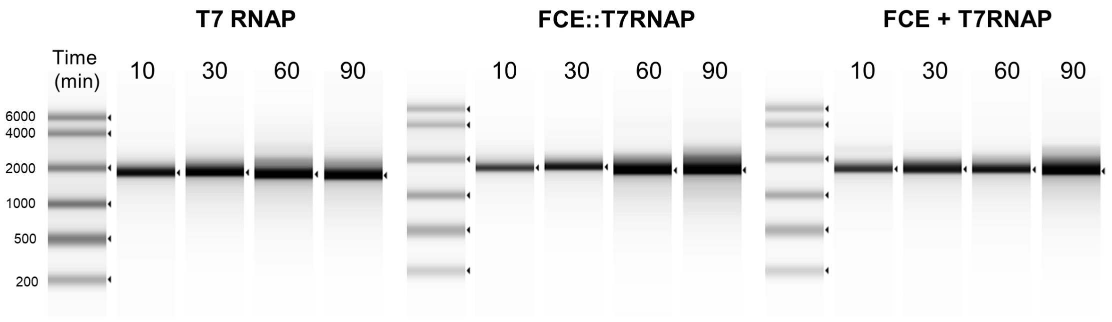

Supporting Data 11

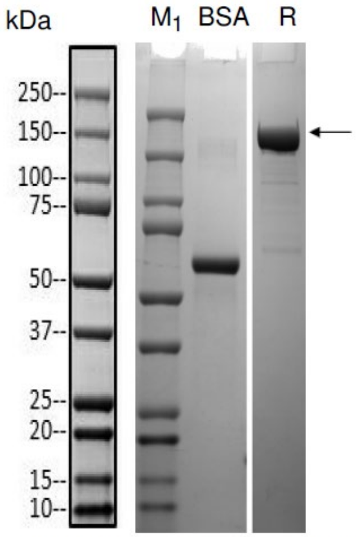


Lane M1: Protein Marker, Bio-rad, Cat. No. 1610374S, refer to annotated key on the left for size
BSA: 2.00 µg
R:Reducing condition

Supporting Data 12

FLuc

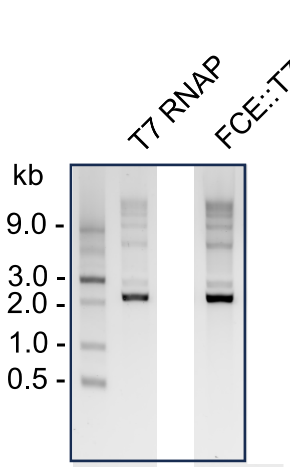


HBB

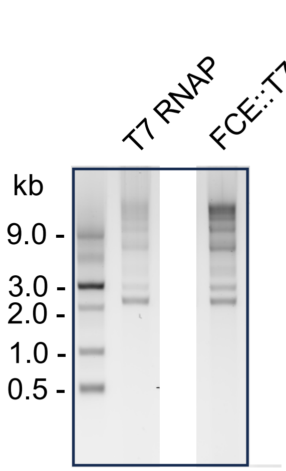


pRNA21

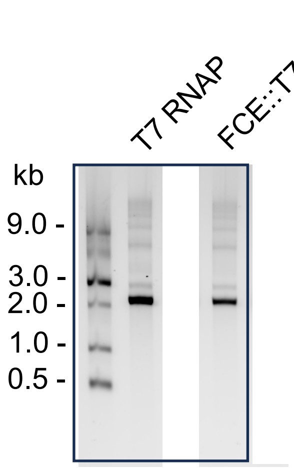


mRNA-1273*

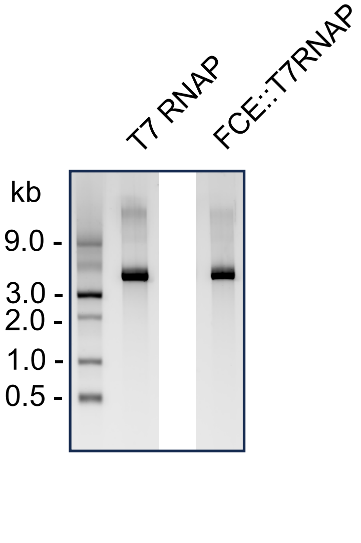
